# Immune Biomarkers, Profiles, and Responses: A Vaccine Ontology Perspective

**DOI:** 10.1101/2025.07.18.665557

**Authors:** Yongqun He, Anthony Huffman, Jie Zheng, Anna Maria Masci, Asiyah Yu Lin, Barry Smith

## Abstract

**Background:** Vaccines have the ability to induce a range of immune responses under different conditions, for example stimulating the production of neutralizing antibodies to block pathogen entry or activating cytotoxic T-cells to eliminate infected cells. Many such immune responses have not been thoroughly examined and classified. The Vaccine Ontology (VO) is a community-based ontology in the domain of vaccinology. We here describe how VO is used to represent the variety of immune responses associated with vaccines, together with associated biomarkers and profiles.

**Results:** The VO differentiates ‘vaccination’ and ‘vaccine immunization.’ The former is a process of administering a vaccine *in vivo*; the latter is the outcome of vaccine induction of immune response. This distinction is critical for understanding both the procedure of vaccination and the resulting immune effects. VO also models and represents various vaccine-induced responses at multiple biological levels, including population, organism, organ/tissue, cell, and gene/protein levels. Such an approach captures the complexity of vaccine-induced immunity, from population-wide trend (for example: herd immunity) to molecular mechanisms. VO defines immune biomarkers as material entities such as neutralizing antibodies that signify humoral immune response, and IFN-gamma that is indicative of cell-mediated responses. Such biomarkers provide measurable indicators of the immune system’s functional state post vaccination, enabling robust evaluation of vaccine efficacy. VO classifies ‘immune response profile’ and ‘correlated profile (or correlate) of immune protection’ as ‘process profiles,’ a class in the Basic Formal Ontology (BFO 2.0). Immune response profiles, such as ‘Th1 (or Th2)- biased profile,’ can be induced by various vaccines and vaccine adjuvants. Different types of ‘correlated profile of immune protection’ are also identified, such as mechanistic and non-mechanistic correlates of immune protection. Such distinctions help us to quickly identify biomarkers and associated prediction and measurement of different kinds of vaccine protection.

**Conclusion:** The important immune-related terms for immune biomarkers, profiles, and responses are modeled ontologically in VO together with their interrelations. The results support enhanced classification and analysis of vaccine-induced immune responses and related biomarkers and immune profiles, leading to further understanding of the vaccine immune mechanisms and enhanced vaccine research and development.

## Background

Infectious diseases remain a major source of human mortality worldwide, contributing to over a quarter of global mortality [1]. Vaccines are biological preparations designed to stimulate the immune system to recognize and fight specific pathogens. As one of the most transformative advances in modern medicine, vaccination has been used to protect not only humans but also cats, dogs, horses, cows, and other animals, against a range of infectious diseases and thereby improve not only human but also animal health. However, our attempts to develop effective and safe vaccines to protect against many diseases have not always been successful [2]. Future success of effective vaccine development will require a thorough understanding of the mechanisms of vaccine-host immune responses and on rational design and development of vaccines with both efficacy and safety.

While vaccines are known to induce adaptive immune responses to infectious pathogens, how specific vaccines induce such immune responses is not always well-understood. For example, there have been many papers discussing how to classify various types of correlates of protection against infection [3–7]. This is an important topic since it is often difficult to evaluate whether a given vaccine candidate can elicit a protective response. The identification of correlates of protection will enable us to better evaluate and predict vaccine levels of protection and thereby support efficient vaccine development and evaluation. An improved understanding not only of correlates of protection but of many related phenomena also faces difficulties. Thus our attention will be directed in what follows also to vaccine-induced responses, vaccine-induced gene responses, vaccine-induced immune correlates of protection, and immune response profiles.

In the era of information-driven biomedicine, an ontology is a human- and computer-understandable representation of types of entities and of relations among such types in the real world, and ontologies so conceived provide a useful platform for the modeling of such immunological topics.

As a biomedical ontology in the domain of vaccines, the Vaccine Ontology (VO) represents the range of different types of vaccines, vaccine formulations, and vaccine responses [8, 9]. VO represents a total of over 10,000 vaccines at different stages (licensed, authorized, in clinical trials, or verified with laboratory animal models). It classifies and defines various vaccines and vaccine components (such as vaccine adjuvants), as well as vaccine induced responses [10]. Underlying functions and immune mechanisms of these vaccine candidates are being further analyzed using ontology-based approaches [11]. In this communication we report how VO represents a number of important factors related to immune response.

## Methods

### General strategy of immune modeling

Immune modeling uses a combination of top-down and bottom-up approaches to ensure comprehensive representation of vaccine immune response. For the top-down approach, we started with the existing VO hierarchy as a foundational framework and added new terms encountered in the literature in alignment with this hierarchy.

For the bottom-up approach, we started out from addressing the issues documented in the VO GitHub web repository issue trackers. For example, a recent focus of our research concerned how to use VO to model correlates of protection against infection, which can be used to predict the levels of protection, where a VO track issue (https://github.com/vaccineontology/VO/issues/603) had been raised. Meanwhile, we have conducted literature surveys and referenced many articles discussing related topics.

### VO ontology development

VO was developed by following the Open Biomedical Ontology (OBO) Foundry ontology development principles, including openness and collaboration [12]. To support ontology interoperability, the development of VO also follows the strategy of “eXtensible Ontology Development” (XOD), which includes four principles: term reuse, semantic alignment, new term addition using ontology design pattern, and community extensibility [13]. The Ontofox tool [14] was used to support the reuse and alignment of terms and relations from existing ontologies. VO imports related terms and relations from other ontologies.

All the terms are aligned under the top-level Basic Formal Ontology (BFO) [12, 15], which is an ISO/IEC ontology standard (ISO/IEC 21838-2:2021). BFO has been adopted by more than 600 ontologies. The usage of BFO allows us to reuse and integrate with ontologies in a way that avoids many interoperability issues and helps make ontologies more widely used as reference vehicles. VO reuses terms from reference ontologies such as the Ontology for Biomedical Investigations (OBI) [16], and Infectious Disease Ontology (IDO) [17]. Ontorat [18] or ROBOT [19] was used to support new VO term generation. The Ontobee [20] was used for ontology term dereferencing.

### VO status, source code, deposition, and license

The VO source code has a CC-BY license and is freely available on the GitHub VO repository (https://github.com/vaccineontology/VO), and VO is available in the Ontobee (https://ontobee.org/ontology/VO), BioPortal (https://bioportal.bioontology.org/ontologies/VO), and OLS repository (https://www.ebi.ac.uk/ols4/ontologies/vo).

## Results

### The upper-level VO structure and design pattern for vaccine immune response modeling

VO reuses and interlinks vaccine-related terms from reference ontologies such as NCBITaxon ontology [21], Gene Ontology (GO) [22], Infectious Disease Ontology (IDO) [17], Disease Ontology (DOID) [23], Ontology for Biomedical Investigations (OBI) [16], and Protein Ontology (PR) [24]. In addition, VO also has generated many new vaccine-specific terms using the method defined in the Methods section.

Figure 1 represents two major branches, for ‘continuant’ and ‘occurrent’, respectively. The ‘continuant’ branch covers entities which exist in time while preserving their identity, such as ‘material entities’ (such as organisms, vaccines, proteins) and ‘realizable entities’ (such as dispositions) that are borne by ‘material entities’ and realized in ‘processes’. Different types of ‘processes’, including ‘vaccination’, ‘immunization’, and ‘response to stimulus’, all fall under the ‘occurrent’ branch. We also deploy the term ‘process profile’ (as in ‘immune response profile’), which is a term originating in BFO. Although not included in the ISO standard version of BFO (BFO 2020), its continued use in many communities led to its inclusion in the mid-level Common Core Ontologies [25], with its original BFO identifier (BFO_0000144).

**Figure 1.**
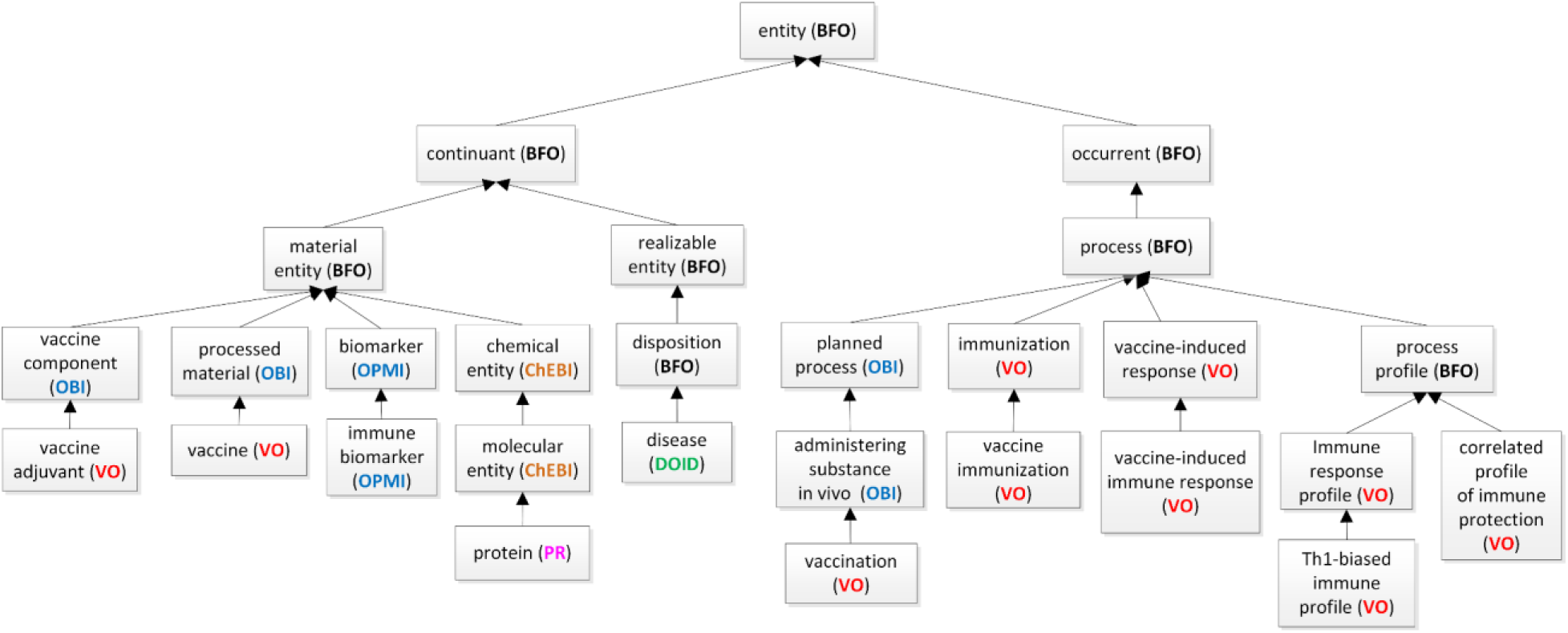
The top-level hierarchical structure of VO as related to immune responses.

In BFO 2.0, ‘process profile’ (BFO_0000144) is defined as follows:

> process profile =*def.* An occurrent that is an occurrent part of some process by virtue of the rate, or pattern, or amplitude of change in an attribute of one or more participants of said process.

Intuitively, a ‘process profile’ is a part of a process which belongs to a dimension of change within that process. In a conversation, for example, the entire process contains many types of subprocesses – for instance the heart beating processes in each of the conversation participants, or the stream of utterances exchanged in the course of the conversation. The latter are process profiles in the BFO intended sense. Often, we are dealing with what we shall call ‘measurement process profiles,’ which are process profiles of a type which can be associated with a time series graph yielded by successive measurements, as for instance in the case of a temperature chart. The pattern exhibited by the rising and falling temperatures in the life of an organism is a measurement process profile, by this definition.

### Ontological modeling of vaccination and immunization

Figure 2 illustrates the differentiates between ‘vaccination’ and ‘vaccine immunization’. The former is a process of administering a vaccine in vivo with the intent to elicit a protective or therapeutic immune response. The latter is a process of priming or modifying a recipient’s adaptive immune response to an antigen. Vaccine immunization thus involves the achieved outcome of vaccine induction of immune response. Vaccination is only the administering process, and some vaccinations do not result in immunization. In contrast, vaccine immunization means the process that leads successfully to the intended outcome.

**Figure 2.**
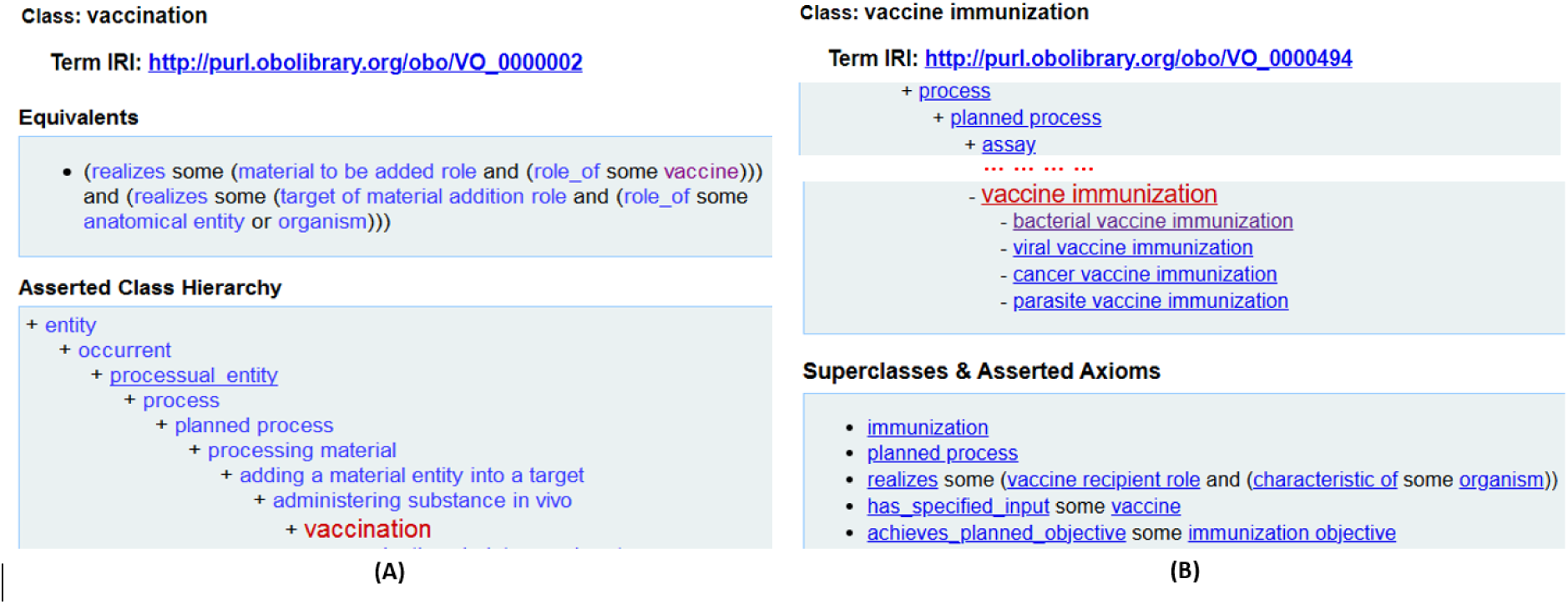
Differentiation of ‘vaccination’ and ‘vaccine immunization’ in VO. (A) Hierarchy of ‘vaccination’ in VO. (B) Hierarchy of ‘vaccination immunization’ in VO.

The following are the formal definitions of ‘vaccination’ (VO_0000002), ‘vaccine immunization’ (VO_0000494), and ‘immunization’ (VO_0000490) in VO:

> vaccination=*def.* Process of administering a vaccine *in vivo* to a recipient (e.g., a human), with the intent to invoke a protective or therapeutic adaptive immune response.

> vaccine immunization = *def.* Immunization that is induced by a vaccine via vaccination process.

> immunization = *def.* Process that results in an adaptive immune response to one or more antigens.

We hereby differentiate the VO: immunization (VO_0000490) and OBI: immunization (OBI_1110094). The term ‘immunization’ (OBI_1110094) is defined in the OBI as the following:

> immunization (OBI_1110094) = *def.* Process of an epitope that is part of or derived from an immunogen coming into contact with adaptive immune cells resulting in these cells acquiring immune effector functions specific for the epitope.

By comparing the two definitions, the VO ‘immunization’ definition offers a broader and more general description. The OBI ‘immunization’ definition provides a more detailed and mechanistic explanation of immunization at the molecular and cellular level. Since the epitope can be part of an antigen or derived from an antigen, there is a slightly different flavor of immunization in OBI versus in VO. However, both refer to a result of adaptive immune response. Therefore, the OBI:immunization is closely related to VO:immunization. We could also conclude that the OBI:immunization is a semantically narrower term than VO:immunization, because OBI:immunization focuses specifically on the interaction between epitopes and adaptive immune cells as the core mechanism of immunization. This is the reason why VO:immunization is used for our vaccine immune modeling research. We have been discussing with the OBI developers for possible agreement and modification.

### VO modeling and representation of vaccine-induced immune response processes

An immune response is a physiological reaction which occurs within an organism to respond to internal factors and exogenous factors such as toxins, viruses, bacteria, or vaccines. There are two major types of immune response: innate and adaptive. The latter includes the humoral immune response, which primarily involves antibody production by B cells, and cell-mediated immunity, which is conducted by T cells that include cytotoxic T cells directly killing infected cells and helper T cells releasing cytokines to help other cell responses.

The Gene Ontology (GO) Biological Process branch has the term ‘immune response’ (GO:0006955), which it defines as “any immune system process that functions in the calibrated response of an organism to a potential internal or invasive threat”. Here as elsewhere the GO deals only with what we can think of as *natural* biological processes. Thus it allows only for natural immune responses [26], where vaccination is a man-made medical intervention and is therefore not within the scope of GO. In addition, GO represents how genes and gene products function only at the molecular level and at the system level, and it is not designed to describe the functions of higher order objects like the heart [26]. This provides the rationale for the inclusion in VO of terms which reference man-made interventions, though we note that many vaccine-induced immune responses can be mapped to GO immune responses, and VO identifies connections to GO immune response terms whenever possible.

VO defines ‘vaccine-induced (host) immune response’ (VO_0000287) as follows:

> ‘vaccine-induced immune response’ (VO_0000287) =*def.* An immune system process that is realized in the calibrated response of the recipient to a vaccine.

As we saw, different types of vaccine-induced responses occur at different levels, including population level (e.g., herd immunity), organism level (e.g., vaccine-induced organism response), system level (e.g., digestive system, immune system), cell level (e.g., B-cell, T-cell), and gene/protein level (e.g., IFN-gamma, IgG antibody). Thus herd immunity is a population level immunity which brings about that a contagious disease has difficulty spreading among a population due to the sufficiently large percentage of people who are immune against the disease. Herd immunity protects an entire community, which includes not only those people who are infected or vaccinated, but also those who are neither. It also provides a degree of protection to those who cannot be vaccinated. All the mentioned levels of vaccine induced immune response are subclasses of the general term ‘vaccine-induced immune response’ (VO_0000287) (Figure 3).

**Figure 3.**
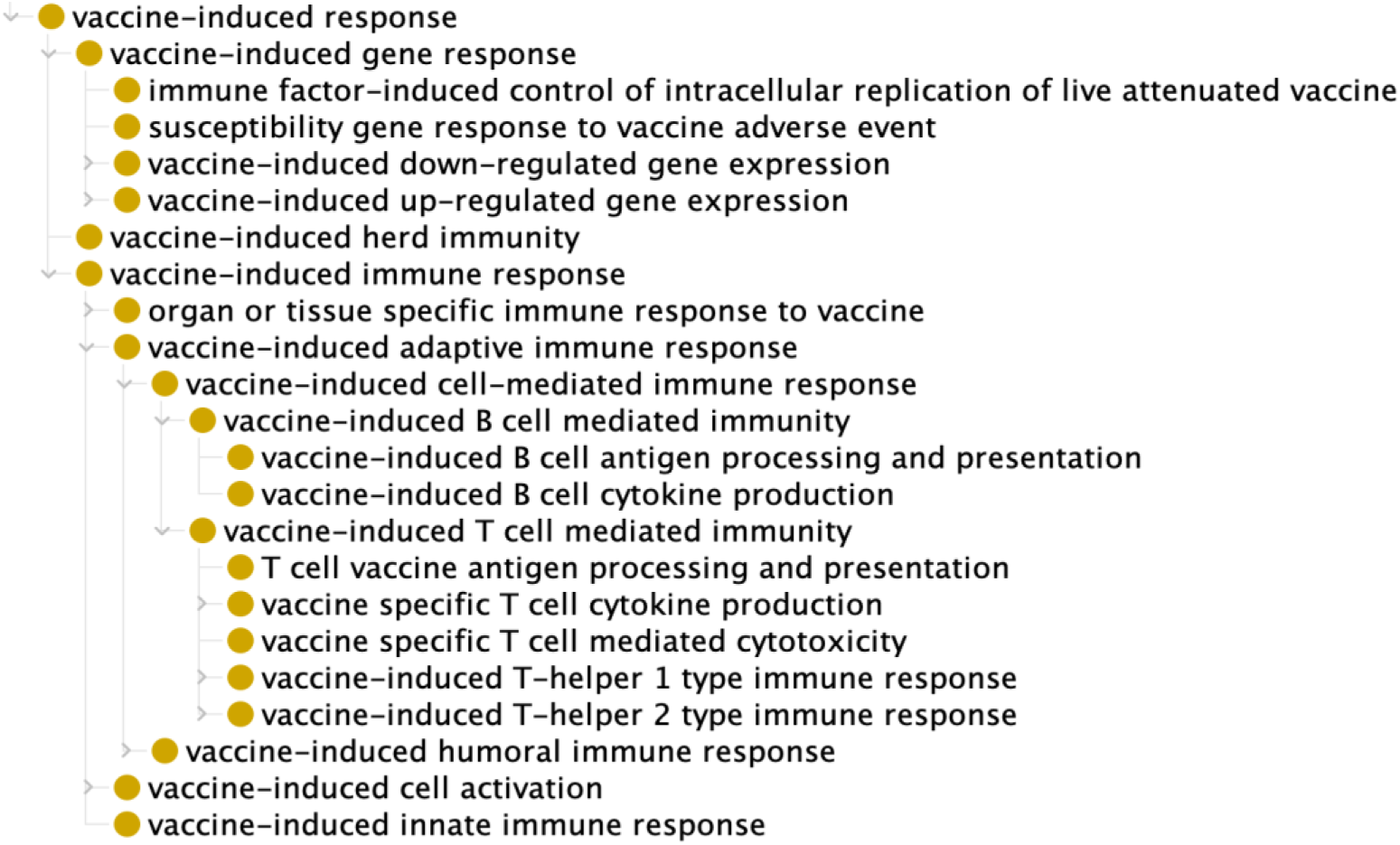
VO modeling of vaccine-induced immune responses.

### VO modeling of biomarkers of vaccine immunity

Two overlapping concepts, namely ‘immune biomarkers’ and ‘correlates of protection*’* (CoP), are often misrepresented in the scientific literature as synonyms. For example, Plotkin and Gilbert indicate that “A CoP is a marker of immune function that statistically correlates with protection after vaccination” [7]. This confusion also makes itself felt in the VO issue tracker (https://github.com/vaccineontology/VO/issues/603). When we examine the literature, however, it becomes clear that the terms ‘immune biomarker’ and ‘correlates of protection’ are used in different ways, so that, for example, modeling different types of correlate of protection helps us to quickly identify the biomarkers that predict the salient level of protection.

VO, therefore, treats these terms as different in meaning. An ‘immune biomarker’ is a material entity (think of it as a material entity present in a certain locus in a host). A ‘correlate of protection’ is a process profile. We first focus on the ontological modeling of immune biomarkers.

A major question is how to define ‘biomarker’ in general. Based on the NIH Biomarkers and Surrogate Endpoint Working Group, a biomarker is defined as “A characteristic that is objectively measured and evaluated as an indicator of normal biological processes, pathogenic processes, or pharmacological responses to a therapeutic intervention” [27]. Unfortunately, the term ‘characteristic’ is itself difficult to define ontologically. Thus, we may say that a certain gene is a biomarker, but then the gene itself is not a ‘characteristic’. Mayeux defined biomarkers as “alterations in the constituents of tissues or body fluids” [28], and here too the term ‘characteristic’ seems inappropriate. Ceusters and Smith proposed a definition according to which there are three disjoint types of biomarker, namely *material*, *biomarker*, and *process biomarkers* [29]. To reap the benefits of defining ‘biomarker’ in such a way that all instances are entities of a single type, we adapt here a proposal advanced by the Ontology of Precision Medicine and Investigation (OPMI) [30, 31] to define biomarker (OPMI_0000445) as a material entity. For our present purposes therefore, we define:

> ‘biomarker’ = *def.* Material entity that has a measurable quality, or that participates in a measurement process profile, which can be used as an indicator of an underlying biological state or identity.

The benefit of defining ‘biomarker’ as a material entity is that no matter what or how we measure, we still need the material entity as the target for the measurement. The material entity is the bearer of the quality or the participant in the process that is measured. This grounding role of material biomarkers makes it possible for us to link to process related biomarkers defined by the NIH Biomarkers and Surrogate Endpoint Working Group.

VO reuses in addition the term ‘immune biomarker’ (OPMI_0000446) from OPMI:

> ‘immune biomarker’ = *def.* Biomarker that has a measurable quality or process profile(s), that can be used as an indicator of an underlying immune state or identity.

Furthermore, VO defines two specific types of immune biomarkers:

> ‘humoral immune biomarker’ (VO_0010463) = *def.* An immune biomarker that indicates a humoral immune response.

and

> ‘cell-mediated immune biomarker’ (VO_0010464) = *def.* An immune biomarker that indicates a cell-mediated immune response.

Humoral immunity is immunity that is mediated by antibodies. Examples of humoral immune biomarkers are: neutralization antibodies, bactericidal antibodies, and IgA biomarkers. For example, the (presence of the) neutralization antibody against COVID-19 correlates statistically with the protection against COVID-19 and so can be considered as a COVID-19 immune biomarker [32–34]. This means that the presence of a neutralization antibody against COVID-19 indicates the induction of protection by a specific COVID-19 vaccine against COVID-19 infection.

The presence of bactericidal antibodies is correlated with the immune protection against *Neisseria meningitidis*, a major cause of meningococcal B (MenB) disease that causes bacterial septicemia and meningitis [35]. The bactericidal antibody biomarker was used in the development of meningococcal B vaccines using the reverse vaccinology strategy [35]. Using *Neisseria meningitidis* serogroup B strain (MC58) genome sequence analysis, Rappuoli *et al.* predicted over 600 positive antigens, of which 350 could be expressed in *E. coli*. These proteins were then purified and administered to immunize separate groups of mice. A total of 91 surface proteins were found to induce antibodies *in vivo*, 29 of which induced antibodies that killed the bacteria *in vitro*, and thereby showed the production of bactericidal antibodies [35]. Even without a virulent challenge assay, therefore, use of the bactericidal antibody biomarker assay allows the rapid identification of many protective antigens. Continuation of this work later resulted in the development of Bexsero, the licensed MenB vaccine [36].

Another group of immune biomarkers are cell-mediated immunity (CMI) biomarkers such as the cytokines Interferon-gamma (IFN-γ) and cytotoxic T lymphocytes (CTL) [37]. IFN-γ is a cytokine that plays an important role in inducing and modulating various immune responses [38, 39]. For example, the *B. abortus* vaccine RB51 is a USDA-licensed cattle vaccine. But an RB51 induced humoral immune response is not protective, as is shown by the inability of the RB51-induced antibody to induce specific protection against virulent *Brucella* infection [37, 40]. On the other hand, RB51 is able to induce specific cell-mediated immunity, as is illustrated by RB51-induced IFN-γ and CTL production [37]. IFN-γ and CTL can therefore be referred to as RB51-induced immune biomarkers whose increased production level correlates with vaccine-induced immune protection.

### VO modeling of immune response profiles

VO classifies the ‘immune response profile’ as a type of ‘process profile,’ which is a subclass of BFO: process (Figure 1). Each instance of the type ‘process profile’ is then also a proper part of some instance of the type ‘process*’.*The ‘immune response profile’ is thus part of the total immune system process behaviour.

VO’s ‘immune response profile’ thus represents a certain recognizable part of the surrounding immune response process, consisting of those interconnected processes that contribute to an immune response. Different types of immune profiles exist corresponding to the components and processes involved. For example, effector T cells have major subtypes, including Type 1 helper T (Th1) cells, Type 2 helper T (Th2) cells, and Type 17 helper T (Th17) cells [41–43]. Each of these T cell subtypes can be induced by specific vaccines or vaccine adjuvants. Correspondingly, we can define different types of immune profiles, including:

> ‘Th1-biased immune profile’ (VO_0005309) =*def.* An immune profile that is characterized by the presence of Type 1 T helper (Th1) cells that produce interferon-gamma, interleukin (IL)-2, and tumor necrosis factor (TNF)-beta, which activate macrophages and are responsible for cell-mediated immunity and phagocyte-dependent protective responses.

> ‘Th2-biased immune profile’ (VO_0005310) =*def.* An immune profile that is characterized by the presence of T type 2 Th (Th2) cells that produce IL-4, IL-5, IL-10, and IL-13, which are responsible for strong antibody production, eosinophil activation, and inhibition of several macrophage functions, thus providing phagocyte-independent protective responses.

> ‘Th17 immune profile’ (VO_0005313) =*def.* An immune profile that is characterized by the presence of T helper 17 cells (Th17), a subset of pro-inflammatory T helper cells defined by their production of interleukin 17 (IL-17). They are related to T regulatory cells and the signals that cause Th17 to differentiate and inhibit Treg differentiation.

And as is shown in Figure 4, there are also mixed immune profiles such as ‘Th1/Th2 mixed immune profile’, ‘Th1/Th17 mixed immune profile’, and ‘Th1/Th2/Th17 mixed immune profile’.

**Figure 4.**
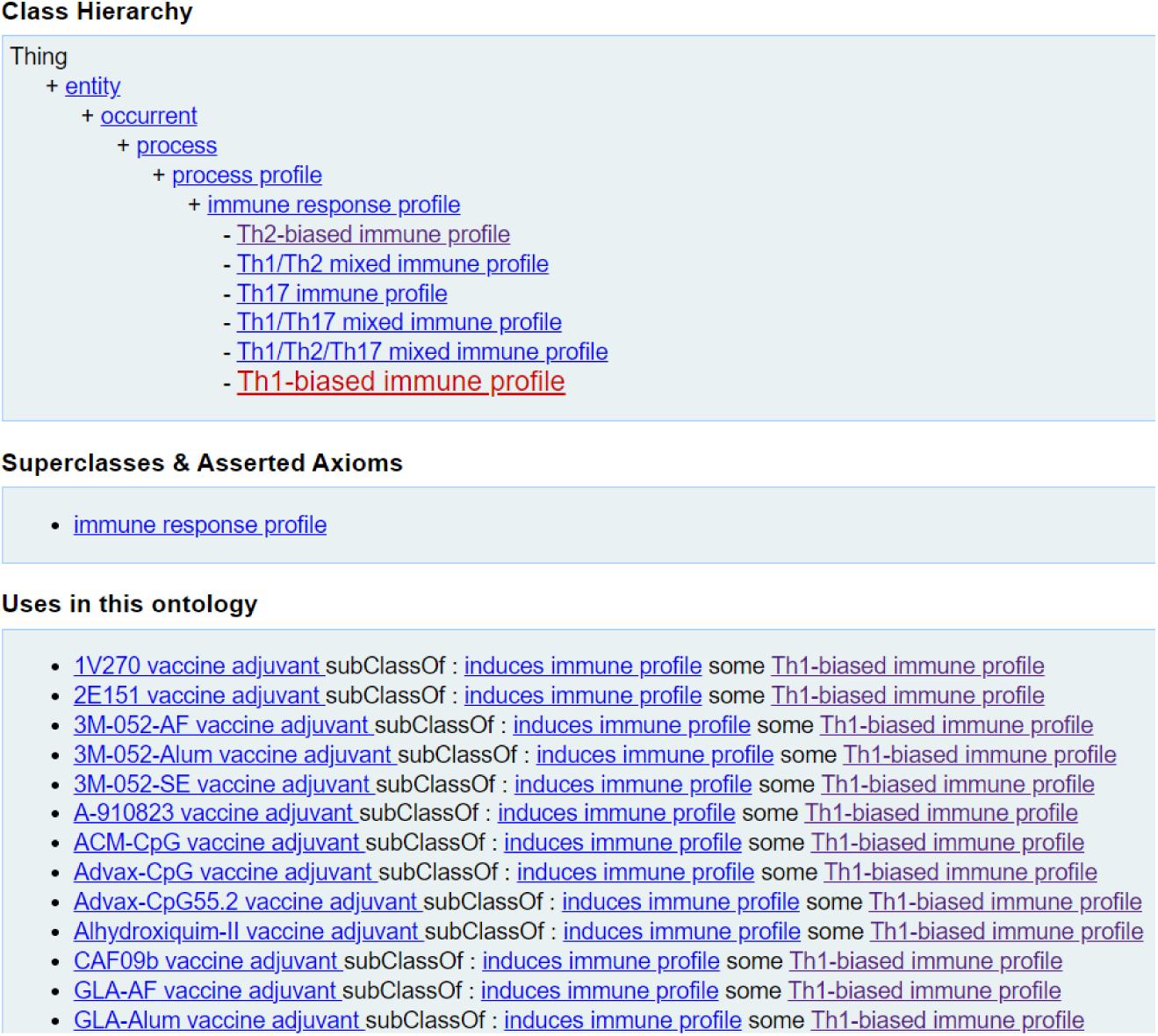
VO classification of ‘Th1-biased immune profile’ and its hierarchy and uses.

Figure 4 shows the classification of the ‘Th1-biased immune profile’ [44], its associated terms, and the relations among them. Specifically, ‘Th1-biased immune profile’ can be induced by different types of vaccine adjuvants such as ‘1V270 vaccine adjuvant’ (VO_0005270) [45] and ‘CpG55.2 vaccine adjuvant’ (VO_0006094) [46]. Note that these and many other vaccine adjuvants represented in the VO are drawn originally from the Vaccine Adjuvant Compendium (VAC) (https://vac.niaid.nih.gov/), an intramural NIH project that collects all the vaccine adjuvants generated through NIH grant support.

### VO modeling of correlated profiles of immune protection

Each immune biomarker (for example each gene or antibody protein) has some expression level or pattern that correlates with protection. We thus refer to the processes – or better to the measurement process profiles – through which these levels or patterns are reached, ‘correlates of protection.’

A correlate of protection is thus not a biomarker, for we have defined biomarkers as material entities. It is, rather, a process profile in BFO terms. Specifically, ‘correlated profile of immune protection’ refers to biological responses that are responsible for and statistically interrelated with protection against a specific infection or disease [6]. Correlates of protection help researchers and clinicians understand what immune mechanisms are most effective in preventing or controlling a given illness. They are often used as indicators of the effectiveness of vaccines or immune-based therapies.

The study of such correlated immune profiles is important because it is typically expensive, time-consuming, and morally irresponsible to perform protection studies directly to specific human cases, and the calling upon identified immune correlates as indicative of immune protection thus saves time by enabling the evaluation of vaccines in a manner that can be applied at a general level. The identification of correlates of protection helps also in designing vaccines that elicit desired immune responses, and correlates can serve also here as surrogate endpoints in clinical trials, thereby accelerating vaccine approval.

In VO, a ‘correlated profile of immune protection’ (with synonym: correlate of immune protection) (VO_0000857) is classified as a BFO:process profile, which can be defined as follows:

> ‘correlated profile of immune protection’ =*def.* Measurement process profile determined by the measurement data generated from a measurement process which indicates the degree of immunological protection in the vaccine recipient.

Here the process profile exists because either a quality or process can be measured in such a way that the measurements form a time series graph. The time series graph then provides an indicator for the degree of immune protection.

As defined above, we can correspondingly define the relation between ‘correlated profile of immune protection’ and ‘immune biomarker,’ using the relation type ‘process_profile_of*’* (BFO_0000133). Specifically, ‘correlated profile of immune protection’ (synonym: ‘correlate of protection’) is a process profile with some immune biomarker as participant:

> ‘correlated profile of immune protection’ =*def.* process_profile_of some (‘vaccine-induced immune response’ and (‘has participant’ some ‘immune biomarker’))

Here the immune biomarker has either a measurable quality or participates in some measurement process profile(s), which in either case can be used as an indicator of an underlying immune protection status. The immune biomarker comes with measurable immune response data that statistically correlates with protection.

To identify such immune biomarkers and their associated correlated immune responses, clinical and laboratory studies are used to examine the relationship between specific immune responses (e.g., antibody titers or T-cell activity) on the one hand and protection from disease on the other. These studies often compare immune responses in vaccinated vs. unvaccinated groups or between individuals who resist infection and those who do not.

Common correlates of immune protection include antibody levels, T-cell responses, and cytokine profiles. High levels of neutralizing antibodies are often a correlate of protection for many viral infections (e.g., influenza, COVID-19). Strong cytotoxic T-cell responses are also correlated with protection, especially in diseases where cellular immunity is critical (e.g., tuberculosis, brucellosis, AIDS, and Epstein-Barr virus infection). Certain cytokines, like interferon-gamma, may correlate with protective immune responses such as *Brucella* vaccine RB51-induced immune protection [37]. Here the interferon-gamma gene expression over-expression induced by RB51 is a correlated profile of immune protection against brucellosis, which is caused by *Brucella*.

Different types of correlates of protection (CoP) [7] are categorized in VO (Figure 5). As detailed below, the categorization is based on a range of criteria, including the causality between CoP and protection, population coverage, and statistical association level.

**Figure 5.**
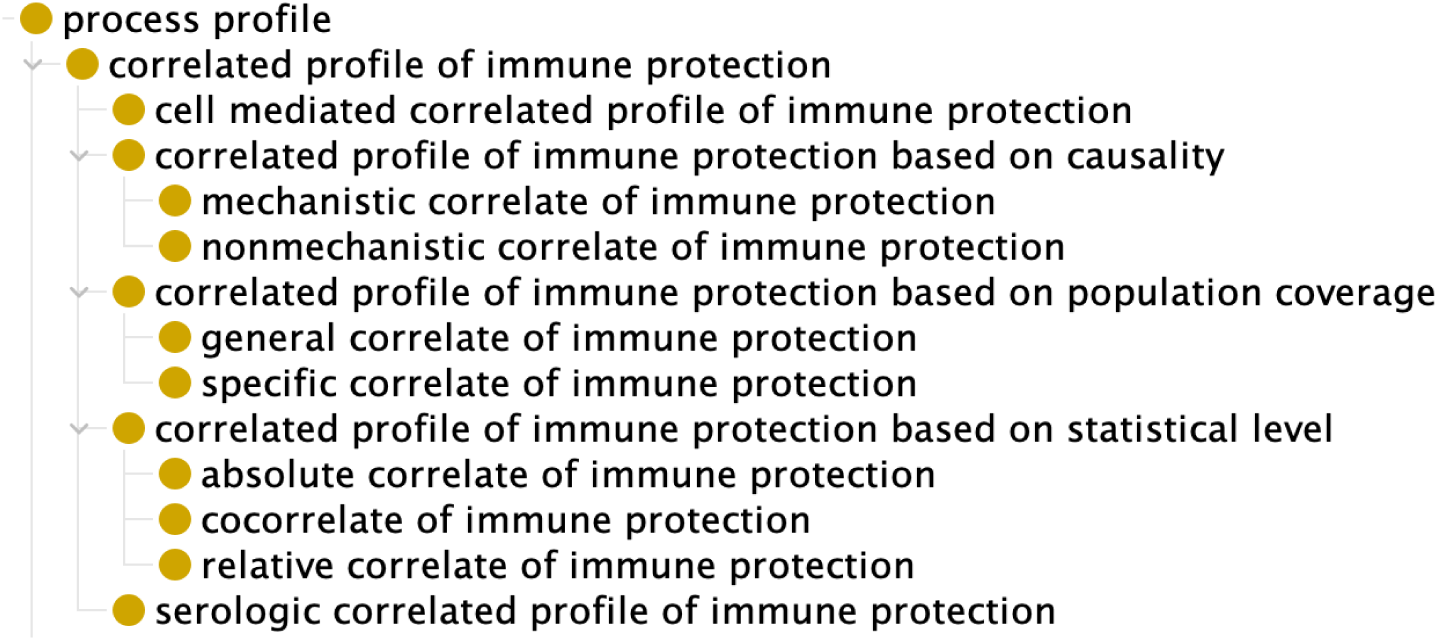
Subtypes of correlated profile of immune protection.

Based on whether there is a causality between the biomarker and the correlated protection, two specific subtypes have been distinguished, namely *mechanistic* and *non-mechanistic* correlates of immune protection (mCoP and nCoP, respectively) [4–7] (Figure 5). An mCoP represents a mechanistic cause of protection. An nCoP does not cause protection; rather, it predicts protection through an identified statistical correlation with an immune response. mCoPs and nCoPs are mutually exclusive. The classification of CoPs, mCoPs, and nCoPs varies with different factors, including the type of vaccine, pathogenesis and pathogenetics of disease, or host population [3, 6]. Examples of mechanistic correlates of protection include neutralization antibody for COVID-19, together with bactericidal antibody humoral immune response biomarkers and CTL cell-mediated immunity biomarkers for brucellosis vaccine. In many cases the neutralizing antibody is an immune marker for the protection against many diseases such as COVID-19 [32–34]. Its profile correlates with protection, it is causing the protection, and it is also a marker of the protection. At the same time, it is often difficult to identify vaccine protective mechanisms. For example, for rotavirus vaccines, a serum immunoglobulin A (IgA) response is elicited in most vaccine recipients [47]. However, serum IgA appears to not be the actual protective mechanism. IgA is then a CoP and an nCoP, but not an mCoP [7].

When CoPs are viewed from the perspective of population coverage, then a distinction is made between specific CoP (or Level 1 CoP) and general CIP (or Level 2 surrogate of protection, or bridging CoP) [7]. Specific CoPs work within a defined population of vaccinees and are predictive of vaccine protection in the same setting as the prior trial. In contrast, general CoPs are predictive of protection in different settings such as across vaccine lots, across human populations, across viral populations, and across species [3].

Based on the statistical level of association, there are absolute correlates of protection (CoPs), relative CoPs, and co-correlates of protection [6]. An absolute CoP has a specific level of response that is highly and positively correlated with protection over a specific threshold. A relative CoP has a level of response that is variably correlated with protection: the correlation varies, it is not always high, and it may depend on various conditions such as population demographics. A co-correlate of protection involves two or more response factors that correlate with protection in alternative, additive, or synergistic ways [6]. For example, cell-mediated immunity may function as a correlate or co-correlate of protection against a disease. When the true correlate of protection is unknown or hard to measure, surrogate tests (e.g., antibody detection) can be used to predict the protection induced by vaccines [5].

## Discussion

This manuscript focuses on using the Vaccine Ontology (VO) to model three important aspects of immunity: immune responses, immune biomarkers, and immune response profiles and their correlated profiles of immune protection. While all of these topics have been well discussed in the immunology literature, we here provide for the first time an ontological perspective that serves to model these topics and the relations between them in a single comprehensive framework.

In contrast to traditional modeling from a pure science perspective, ontology modeling is based on a logical approach (using the OWL Description Logic), which focuses on transforming scientific understanding into a hierarchical taxonomic framework rooted in the ‘is a’ (for: *is a subtype of*) relation, together with a treatment of other relations, such as ‘part of’ and ‘process profile of,’ to further represent the relations among types of entities in the taxonomy. In addition, the ontology terms and relations are both human- and computer-understandable, making the ontological modeling and representation suitable for the modeling of the complex immune related entities and relations, resulting in computational applications of these modeling results (increasingly including applications in AI). The Vaccine Ontology (VO) modeling described in this communication demonstrates clearly how definitions of terms and formulation of axioms enable deep ontological modeling of a sort that can yield a better understanding of complex biological phenomena and can also be used in applications for example pertaining to regulation and innovation in the vaccine domain.

Our model illustrates how the three types of immune response processes: vaccine-induced immune response, immune response profile, and correlated profile of immune protection are closely related. An immune response profile represents a pattern of immune response that encompasses various interconnected processes and components of the immune system. Examples – such as Type 1 helper T (Th1) or Th2 immune profile – represent a type of immune response induced by specific types or groups of vaccines or vaccine adjuvants. A correlated profile differs from an immune response profile in that it correlates with immune protection, and it may not be identical to or be a part of the immune protection itself.

The modeling of such phenomena is challenging and terms are often used in a confused way, for example because it uses ‘vaccination’ and ‘immunization’ interchangeably. However, our modeling distinguishes one from the other based on whether an immune response is or is not elicited. The purpose of vaccination is to establish protective immune responses. But the successful establishment of an immune response depends on many interconnected factors. There are all kinds of immune responses at different levels, and these may be summarized as immune response profiles. To deeply understand the complex immune mechanisms involved, it is critical to work with a systematic classification based on a taxonomy whose terms are rigorously defined and in such a way as to capture the relations between them. The immune biomarkers and correlates of immune protection we have focused on here are also often handled in a confused fashion by many, for example because correlations of protection are measured via levels of the biomarker only for one specific disease. One novel contribution of this study is that we systematically separate out such factors. In this way we can recognize that biomarkers and correlates or protection are different but at the same time closely related. Not all diseases have clear or singular correlates of protection. In some cases, multiple immune mechanisms work together to provide protection. Also, statistical correlations do not always imply causation. Understanding different types of correlations is fundamental in immunology and translational medicine since it bridges the gap between immune response mechanisms and real-world disease prevention.

Using the ontological immune modeling as the overall framework, we have also modeled and represented specific subtypes of immune responses, profiles, and biomarkers. For example, we have represented vaccine-induced immune responses at the different levels, including population, organism, organ, tissue, cell, and molecular gene and protein levels. We have certainly not addressed all possible classification of all the responses at these different levels.

The complexity of the interactions among the tissue, immunity, cells, and signals at different conditions [48] requires careful examination and modeling. However, we believe that the ontological framework we have laid out paves the way for further in-depth study and ontological modeling for more subtypes or subcategories of these important immune entities.

Note that the modeling results represent not only the views of our VO developer team, but also of many vaccine researchers and of ontologists working in neighboring fields. They also represent the results of our communications with the vaccine community as recorded in the VO tracker (https://github.com/vaccineontology/VO/issues/603). We realize that our terminological choices do not always align with the vaccine community in its entirety – not least in those areas where the community itself is divided. However, we welcome discussions with members of this community that will help us to achieve terminological consistency.

Since ontology is based on Description Logic, many ontology-based AI applications are being developed. For example, we can develop Description Logic (DL) queries or SPARQL (i.e., "SPARQL Protocol and RDF Query Language") queries to query the ontology information [20]. We can also develop ontology-based natural language processing (NLP) methods to support efficient extraction of immune related terms from the literature [8, 49–52].

## List of Abbreviations

BFO: Basic Formal Ontology
GO: Gene Ontology
IDO: Infectious Disease Ontology
OBI: Ontology for Biomedical Investigations
OBO: The Open Biological and Biomedical Ontologies
OWL: Web Ontology Language
RDF: Resource Description Framework
SPARQL: SPARQL Protocol and RDF Query Language
UBERON: Uberon multi-species anatomy ontology
VO: Vaccine Ontology

## Declarations

### Human Ethics and Declarations Sections

Not applicable.

### Consent to Publish declaration

Not applicable.

### Availability of data and material

Related data, including the VO source code, is freely available on the GitHub website https://github.com/vaccineontology/VO.

### Authors’ Contributions

YH: VO developer, vaccine and immune response domain expert, project design, and the first version manuscript preparation. AH and JZ: VO developer, collection and modeling of vaccines and vaccine formulations. JZ, AMM and AYL: Immune response modeling. BS: BFO developer ensuring VO alignment with BFO, ontological modeling and consultation. All authors contributed to manuscript discussion and preparation.

## Acknowledgements

We acknowledge the discussion with experts in the vaccine and immunology communities. Part of the work was initially presented in the 2023 Workshop of Ontologies for Infectious and Immune-Mediated Disease Data Science (OIIMDDS 2023) workshop during the 2023 International Conference on Biomedical Ontology (ICBO 2023, https://delaneycdmcnulty.wixsite.com/oiids-workshop), University of Brasilia, Brazil. We appreciate the submission of a related VO issue tracker by Ms. Lindsey N Anderson from Pacific Northwest National Laboratory (PNNL), and the following discussion by Mr. Jeremy Zucker from PNNL and Mr. Charles Tapley Hoyt from the OBO Foundry Operation Team.

## References

1. Becker K, Hu Y, Biller-Andorno N: Infectious diseases - a global challenge. Int J Med Microbiol 2006, 296(4-5):179–185.

2. Hadfield J, Megill C, Bell SM, Huddleston J, Potter B, Callender C, Sagulenko P, Bedford T, Neher RA: Nextstrain: real-time tracking of pathogen evolution. Bioinformatics 2018, 34(23):4121–4123.

3. Qin L, Gilbert PB, Corey L, McElrath MJ, Self SG: A framework for assessing immunological correlates of protection in vaccine trials. J Infect Dis 2007, 196(9):1304–1312.

4. Plotkin SA: Immunologic correlates of protection induced by vaccination. Pediatr Infect Dis J 2001, 20(1):63–75.

5. Plotkin SA: Vaccines: correlates of vaccine-induced immunity. Clin Infect Dis 2008, 47(3):401–409.

6. Plotkin SA: Correlates of protection induced by vaccination. Clin Vaccine Immunol 2010, 17(7):1055–1065.

7. Plotkin SA, Gilbert PB: Nomenclature for immune correlates of protection after vaccination. Clin Infect Dis 2012, 54(11):1615–1617.

8. Ozgur A, Xiang Z, Radev DR, He Y: Mining of vaccine-associated IFN-gamma gene interaction networks using the Vaccine Ontology. J Biomed Semantics 2011, 2 **Suppl 2**(Suppl 2):S8.

9. Lin Y, He Y: Ontology representation and analysis of vaccine formulation and administration and their effects on vaccine immune responses. J Biomed Semantics 2012, 3(1):17.

10. Huang PC, Goru R, Huffman A, Yu Lin A, Cooke MF, He Y: Cov19VaxKB: A Web-based Integrative COVID-19 Vaccine Knowledge Base. Vaccine X 2021:100139.

11. Huffman A, Masci AM, Zheng J, Sanati N, Brunson T, Wu G, He Y: CIDO ontology updates and secondary analysis of host responses to COVID-19 infection based on ImmPort reports and literature. J Biomed Semantics 2021, 12(1):18.

12. Smith B, Ashburner M, Rosse C, Bard J, Bug W, Ceusters W, Goldberg LJ, Eilbeck K, Ireland A, Mungall CJ et al: The OBO Foundry: coordinated evolution of ontologies to support biomedical data integration. Nat Biotechnol 2007, 25(11):1251–1255.

13. He Y, Xiang Z, Zheng J, Lin Y, Overton JA, Ong E: The eXtensible ontology development (XOD) principles and tool implementation to support ontology interoperability. J Biomed Semantics 2018, 9(1):3.

14. Xiang Z, Courtot M, Brinkman RR, Ruttenberg A, He Y: OntoFox: web-based support for ontology reuse. BMC research notes 2010, 3:175:1–12.

15. Arp R, Smith B, Spear AD: Building Ontologies with Basic Formal Ontology. MIT Press: Cambridge, MA, USA; 2015.

16. Bandrowski A, Brinkman R, Brochhausen M, Brush MH, Bug B, Chibucos MC, Clancy K, Courtot M, Derom D, Dumontier M et al: The Ontology for Biomedical Investigations. PLoS One 2016, 11(4):e0154556.

17. Babcock S, Beverley J, Cowell LG, Smith B: The Infectious Disease Ontology in the age of COVID-19. J Biomed Semantics 2021, 12(1):13.

18. Xiang Z, Zheng J, Lin Y, He Y: Ontorat: automatic generation of new ontology terms, annotations, and axioms based on ontology design patterns. J Biomed Semantics 2015, 6:4.

19. Jackson RC, Balhoff JP, Douglass E, Harris NL, Mungall CJ, Overton JA: ROBOT: A Tool for Automating Ontology Workflows. BMC Bioinformatics 2019, 20(1):407.

20. Ong E, Xiang Z, Zhao B, Liu Y, Lin Y, Zheng J, Mungall C, Courtot M, Ruttenberg A, He Y: Ontobee: A linked ontology data server to support ontology term dereferencing, linkage, query and integration. Nucleic Acids Res 2017, 45(D1):D347–D352.

21. Federhen S: The NCBI Taxonomy database. Nucleic Acids Res 2012, 40(Database issue):D136–143.

22. Ashburner M, Ball CA, Blake JA, Botstein D, Butler H, Cherry JM, Davis AP, Dolinski K, Dwight SS, Eppig JT et al: Gene ontology: tool for the unification of biology. The Gene Ontology Consortium. Nat Genet 2000, 25(1):25–29.

23. Kibbe WA, Arze C, Felix V, Mitraka E, Bolton E, Fu G, Mungall CJ, Binder JX, Malone J, Vasant D et al: Disease Ontology 2015 update: an expanded and updated database of human diseases for linking biomedical knowledge through disease data. Nucleic Acids Res 2015, 43(Database issue):D1071–1078.

24. Natale DA, Arighi CN, Blake JA, Bona J, Chen C, Chen SC, Christie KR, Cowart J, D’Eustachio P, Diehl AD et al: Protein Ontology (PRO): enhancing and scaling up the representation of protein entities. Nucleic Acids Res 2017, 45(D1):D339–D346.

25. Jensen M, De Colle G, Kindya S, More C, Cox AP, Beverley J: The Common Core Ontologies. arXiv preprint arXiv:240417758 2024.

26. Smith B: Biomedical ontologies. In: Terminology, Ontology and their Implementations. Springer; 2023: 125–169.

27. Biomarkers Definitions Working G: Biomarkers and surrogate endpoints: preferred definitions and conceptual framework. Clin Pharmacol Ther 2001, 69(3):89–95.

28. Mayeux R: Biomarkers: potential uses and limitations. NeuroRx 2004, 1(2):182–188.

29. Ceusters W, Smith B: Biomarkers in the ontology for general medical science. Stud Health Technol Inform 2015, 210:155–159.

30. He Y, Ong E, Schaub J, Dowd F, O’Toole JF, Siapos A, Reich CG, Seager S, Wang L, Yu H: OPMI: the Ontology of Precision Medicine and Investigation and its Support for Clinical Data and Metadata Representation and Analysis. In: ICBO: 2019. 1–10.

31. Ong E, Wang LL, Schaub J, O’Toole JF, Steck B, Rosenberg AZ, Dowd F, Hansen J, Barisoni L, Jain S et al: Modelling kidney disease using ontology: insights from the Kidney Precision Medicine Project. Nat Rev Nephrol 2020, 16(11):686–696.

32. Li C, Yu J, Issa R, Wang L, Ning M, Yin S, Li J, Wu C, Chen Y: CoronaVac-induced antibodies that facilitate Fc-mediated neutrophil phagocytosis track with COVID-19 disease resolution. Emerg Microbes Infect 2024:2434567.

33. Bratcher A, Kao SY, Chun K, Petropoulos CJ, Gundlapalli AV, Jones J, Clarke KEN: Quantitative SARS-CoV-2 Spike Receptor-Binding Domain and Neutralizing Antibody Titers in Previously Infected Persons, United States, January 2021-February 2022. Emerg Infect Dis 2024, 30(11):2352–2361.

34. Yang J, Huo X, Jiang Q, Liao Y, Zhang C, Yu L, Wang Q, Niu T, Li C, Pi N et al: Preclinical safety evaluation of intradermal SARS-CoV-2 inactivated vaccine (Vero cells) administration in macaques. Vaccine 2023, 41(17):2837–2845.

35. Pizza M, Scarlato V, Masignani V, Giuliani MM, Arico B, Comanducci M, Jennings GT, Baldi L, Bartolini E, Capecchi B et al: Identification of vaccine candidates against serogroup B meningococcus by whole-genome sequencing. Science 2000, 287(5459):1816–1820.

36. Vernikos G, Medini D: Bexsero(R) chronicle. Pathog Glob Health 2014, 108(7):305–316.

37. He Y, Vemulapalli R, Zeytun A, Schurig GG: Induction of specific cytotoxic lymphocytes in mice vaccinated with Brucella abortus RB51. Infect Immun 2001, 69(9):5502–5508.

38. Schoenborn JR, Wilson CB: Regulation of interferon-gamma during innate and adaptive immune responses. Adv Immunol 2007, 96:41–101.

39. Alspach E, Lussier DM, Schreiber RD: Interferon gamma and Its Important Roles in Promoting and Inhibiting Spontaneous and Therapeutic Cancer Immunity. Cold Spring Harb Perspect Biol 2019, 11(3).

40. Jimenez de Bagues MP, Elzer PH, Jones SM, Blasco JM, Enright FM, Schurig GG, Winter AJ: Vaccination with Brucella abortus rough mutant RB51 protects BALB/c mice against virulent strains of Brucella abortus, Brucella melitensis, and Brucella ovis. Infect Immun 1994, 62(11):4990–4996.

41. Korn T, Oukka M, Kuchroo V, Bettelli E: Th17 cells: effector T cells with inflammatory properties. Semin Immunol 2007, 19(6):362–371.

42. Supriya R, Gao Y, Gu Y, Baker JS: Role of Exercise Intensity on Th1/Th2 Immune Modulations During the COVID-19 Pandemic. Front Immunol 2021, 12:761382.

43. Pulendran B: Modulating TH1/TH2 responses with microbes, dendritic cells, and pathogen recognition receptors. Immunol Res 2004, 29(1-3):187–196.

44. Berger A: Th1 and Th2 responses: what are they? BMJ 2000, 321(7258):424.

45. Wu CC, Crain B, Yao S, Sabet M, Lao FS, Tawatao RI, Chan M, Smee DF, Julander JG, Cottam HB et al: Innate immune protection against infectious diseases by pulmonary administration of a phospholipid-conjugated TLR7 ligand. J Innate Immun 2014, 6(3):315–324.

46. Li L, Honda-Okubo Y, Huang Y, Jang H, Carlock MA, Baldwin J, Piplani S, Bebin-Blackwell AG, Forgacs D, Sakamoto K et al: Immunisation of ferrets and mice with recombinant SARS-CoV-2 spike protein formulated with Advax-SM adjuvant protects against COVID-19 infection. Vaccine 2021, 39(40):5940–5953.

47. Franco MA, Angel J, Greenberg HB: Immunity and correlates of protection for rotavirus vaccines. Vaccine 2006, 24(15):2718–2731.

48. Matzinger P: Friendly and dangerous signals: is the tissue in control? Nat Immunol 2007, 8(1):11–13.

49. Ozgur A, Hur J, He Y: The Interaction Network Ontology-supported modeling and mining of complex interactions represented with multiple keywords in biomedical literature. BioData Min 2016, 9:41.

50. Huffman A, Ong E, Hur J, D’Mello A, Tettelin H, He Y: COVID-19 vaccine design using reverse and structural vaccinology, ontology-based literature mining and machine learning. Brief Bioinform 2022, 23(4).

51. He Y, Yu H, Ong E, Wang Y, Liu Y, Huffman A, Huang HH, Beverley J, Hur J, Yang X et al: CIDO, a community-based ontology for coronavirus disease knowledge and data integration, sharing, and analysis. Sci Data 2020, 7(1):181.

52. He Y, Yu H, Huffman A, Lin AY, Natale DA, Beverley J, Zheng L, Perl Y, Wang Z, Liu Y et al: A comprehensive update on CIDO: the community-based coronavirus infectious disease ontology. J Biomed Semantics 2022, 13(1):25.

